# pH induced motility pattern change in a marine dinoflagellate

**DOI:** 10.1101/2025.07.29.667351

**Authors:** Arshad Kalliyil, Shri Gowri R. Tikoti, S Anudarsh, Sougata Roy

## Abstract

Flagella-driven motility is a conserved feature across eukaryotic lineages, from unicellular plankton to mammals. In marine dinoflagellates, such as *Lingulodinium polyedra*, motility underlies diel vertical migration (DVM), a key adaptive strategy that enables access to spatio-temporally segregated resources in the water column. To investigate how pH influences motility, we used *L. polyedra* and two other dinoflagellates as a model and used a multi-particle tracking algorithm to monitor and quantitatively analyze cellular motility. Under normal pH conditions, *L. polyedra* displays linear, random motility with variable speeds. Upon CO_²_-induced acidification, we observed a dose-dependent decrease in motility speed accompanied by a striking behavioral shift: within minutes of pH reduction, over 90% of the cells transitioned to spiral motility. This effect was both reversible and reproducible, including when pH was modified chemically rather than via CO_²_. The rapid onset of these changes suggests a non-genomic mode of regulation, an area that remains largely unexplored in phytoplankton. We hypothesize that external pH modulates flagellar dynamics in dinoflagellates. Our findings offer new insights into the link between environmental pH and flagellar motility and provide a platform for investigating non-genomic responses to ocean acidification in marine phytoplankton.

---

Motility in marine dinoflagellates ensures optimization of resource acquisition, which is crucial for their survival. Tumbling, swirling and vertical migration of motile dinoflagellates is important for phototaxis, nutrient uptake, photosynthesis, bloom formation [1] and escape predation [2, 3]. Eukaryotic flagellar components that enable cellular motility are extremely conserved across species [4]. Factors such as changes in intracellular calcium levels [5] and the regulation of post-translational modifications (PTMs) of tubulin [6] can disrupt normal flagella-driven motility.

Bacterial and eukaryotic flagella both mediate cell motility but differ markedly in their structural organization and mechanisms of action. In freshwater green algae, prolonged acidification has been shown to reduce phototactic responses by downregulating genes associated with photoreception, signal transduction, and flagellar motility [7]. In contrast, bacterial motility relies on proton-driven, pH-sensitive flagellar motors, resulting in diminished movement under acidic conditions [8]. While pH regulation of phytoplankton motility has largely been attributed to genomic and transcriptional responses, the contribution of non-genomic mechanisms remains largely unexplored. In this study, we did minor modifications in the multi-particle tracking algorithm (Trackpy) to quantitatively assess the impact of pH on marine phytoplankton motility. Our findings provide new insights into how acidified environments influence phytoplankton behavior, offering a broader understanding of cellular responses to environmental stress. Harmful algal blooms (HABs) can induce significant pH shifts in the water column, particularly under confined conditions, and it is plausible that nighttime pH levels during such events may drop to levels that mimic acidified environments that we have shown to affect cellular motility. However, the ecological significance of these pH-induced changes in swimming speed and motility patterns remains to be elucidated.

We used motile cells of *Lingulodinium polyedra*, a representative photosynthetic marine dinoflagellate known for exhibiting robust daily rhythms in vertical migration. To analyze their motility, we employed Trackpy package (v0.6.2) (see Methods for details). Using this tool, we quantitatively assessed the movement patterns and swimming speeds of individual *L. polyedra* cells cultured under standard conditions (pH 8.5 ± 0.2). Under these conditions, the average swimming speed was approximately 136 µm·s^−1^, displaying characteristics typical of a random walk (Figure 1A). The spatial distribution of instantaneous speeds revealed that *L. polyedra* cells can reach peak velocities of up to 500 µm·s^−1^, which intermittently drop to 0 µm·s^−1^ when the cells pause or change direction (Figure 1A, right panel).

**Figure 1:**
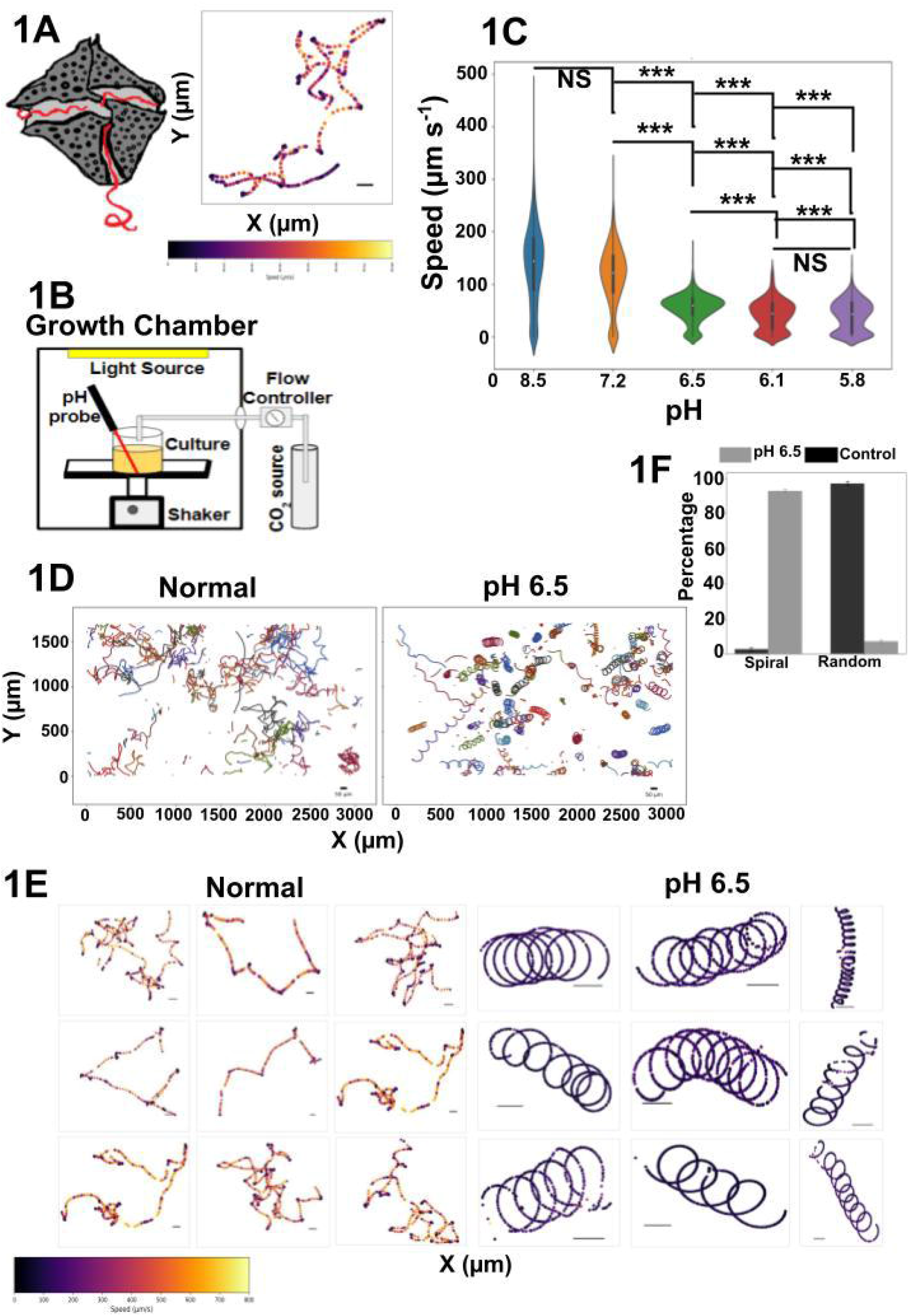
pH induced cellular motility changes. (1A) Left panel: The schematics of a single *L. polyedra* cell with two flagella, one longitudinal and another transverse. Right Panel: The beating of two flagella result in the random motility pattern captured with a Leica imaging system at 4X magnitude for about 22 secs. The captured image was processed with Trackpy package to obtain the motility pattern. The color map of trajectory was plotted using matplotlib in python. X and Y-axis are in µm, and the scale bar = 50 µm. (1B) The schematics describing the installation of the CO_2_ unit to manipulate the medium with desired amount of CO_2_. The pH monitoring and maintenance of the *L. polyedra* cell in pH modified and normal medium. (1C) Dose dependent speed change in motility. Violin plots representing the speed of cells in normal and pH-modified medium was generated using the seaborn package in Python. The non-parametric Kruskal-Wallis Test was done to determine the statistical significance between groups. The p-Value was 1.5e-181. Once we ensure that groups are significantly different, we performed the Post Hoc Dunn Test to find the significance between the groups. The (^***^) signifies P<0.001, and NS denotes not significant. The pattern is bimodal. (1D) Motility patterns either in normal or acidic pH. The top panels shows the cell motility pattern of all tracked cells from a 30 second video. (1E) The color map of the spatial speed distribution pattern of 9 individual cells under normal and acidic pH chosen from all 3 different biological replicates. The scale bar = 50 µm. (1F) The histogram showing percentage of random or spiral motile cells under normal or acidic pH in 3 different independent replicates. The error bar denotes SEM.

To simulate ocean microclimate conditions, we used an algal growth chamber (Percival) equipped with a CO_²_ delivery system designed to regulate pH levels in the culture medium (Figure 1B). Cell motility was assessed 5–20 minutes following each treatment. We observed a significant, dose-dependent decrease in swimming speed in response to acidification (Figure 1C). While similar deceleration has been reported in green phytoplankton following prolonged exposure to low pH conditions [7], our findings demonstrate that such effects can manifest within minutes. Interestingly, we also observed a distinct shift in motility behavior. Upon 5–20 minutes of exposure to acidic medium, *L. polyedra* cells transitioned from random swimming to a consistent spiral trajectory (Figure 1D; Supplementary Video S1). Spatial speed distribution analysis revealed that in acidic conditions, individual cells exhibited slow but uniform spiral motion, a striking contrast to the variable speeds and direction changes seen under normal pH (Figure 1E). This spiral swimming pattern closely resembles the motility behavior reported for *Effrenium voratum* cultured under non-acidified conditions [9]. Across three independent biological replicates, we found that more than 90% of the cells adopted this spiral and decelerated motility pattern when exposed to low pH (Figure 1F).

The pH-induced shift in motility from random to spiral exhibits a clear dose-dependent response. As the external pH gradually decreases, the proportion of cells exhibiting spiral motility increases, reaching a maximum at pH 6.5. Further acidification beyond this point leads to a progressive decline in both spiral and random motility (Figure 2A). In contrast, the percentage of non-motile cells remains relatively constant across a range of pH levels until pH 6.5, after which there is a sharp increase, likely due to acid shock-induced flagellar shedding (Figure 2A). To assess the reversibility of the motility response, we exposed cells to pH 6.5 for 10 minutes and then restored the medium to normal pH (8.5). As hypothesized, over 90% of the cells reverted to random motility after pH normalization (Figure 2B), indicating that the spiral motility phenotype is transient and pH-dependent. To rule out the possibility that changes in gas composition, rather than pH, drive the observed motility shift, we used nitrogen gas (N_²_) at a rate five times higher than that used for CO_²_. As expected, neither the pH nor the motility speed and pattern were affected, confirming that inert gas perturbation alone does not influence cell behavior (Figure 2C). Finally, to directly test the role of pH independent of CO_²_, we chemically adjusted the medium to achieve pH levels comparable to those induced by CO_²_ enrichment. The resulting motility changes mirrored those observed under CO_²_-induced acidification, further confirming that pH, rather than CO_²_ per se, is the primary driver of the motility phenotype (Figure 2C). Harmful algal blooms (HABs) can induce significant pH shifts in the water column [10], particularly under confined conditions, and it is plausible that nighttime pH levels during such events may drop to levels that mimic acidified environments that we have shown to affect cellular motility. However, the ecological significance of these pH-induced changes in swimming speed and motility patterns remains to be elucidated.

**Figure 2:**
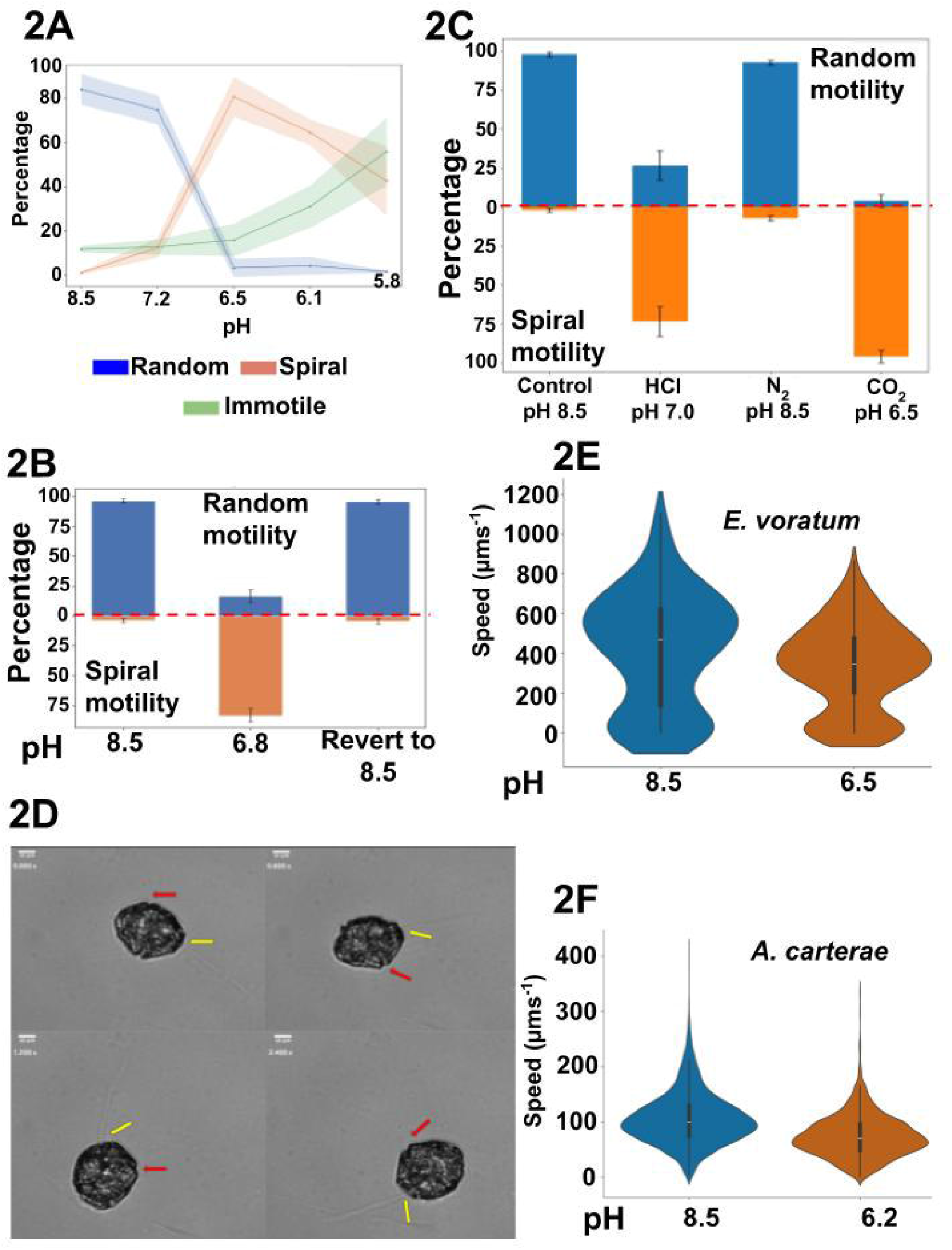
pH induced motility pattern is reversible and not due to asynchronous loss of flagella. (2A) The histogram bar plot shows the percentage of random (blue) or spiral (orange) motile cells under normal pH, acidic pH (induced by HCl), N_2_ treated medium or acidic pH (induced by CO_2_). (2B) Dose dependent percentage of motility pattern change corresponding to the change in pH on X-axis and percent motility on Y-axis. The shaded region represents SEM. (2C) pH reversal reverts the motility pattern. The pH is represented on the X-axis while the Y-axis represents the % motile cells. (2D) The captured video used to visualize the flagella was processed into a composite image using Fiji. The image depicts a single *L. polyedra* cell undergoing spiral motion, with the red and yellow arrows indicating the two distinct flagella. pH-induced changes in swimming speed were observed in (2E) *E. voratum* and (2F) *A. carterae*. Each condition was tested in at least three biological replicates across three different days, with all experiments conducted at the same time of day to control for potential circadian effects. To assess the distribution of the data, the Shapiro–Wilk test was performed. Since the data did not follow a normal distribution, a non-parametric Mann–Whitney U test was used to determine statistical significance. The result was significant, with a p-value < 0.001.

We investigated whether asynchronous flagellar shedding contributes to the altered motility patterns observed in *L. polyedra*. Using polarized light microscopy, equipped with an ultrafast frame imaging system revealed that both flagella remained intact in cells exhibiting spiral motion under acidified conditions (Figure 2D and Supplementary figure S1). Expanding our investigation, we examined the effect of acidification on the motility of *E. voratum* (Figure 2E) and *Amphidinium carterae* (Figure 2F). We observed that while acidification significantly decreased the motility speed of both these dinoflagellates, their distinct swimming patterns were unaffected. Although the precise mechanism underlying the changes in motility speed and pattern remains unclear, the rapid transition of speed and pattern from random to spiral movement within minutes of pH alteration suggests a non-genomic mode of regulation. We believe that acid-induced flagellar dysfunction may alter its beat frequency and waveform, driving the shift in motility behavior. Given the evolutionary conservation of tubulin and its modifying enzymes in dinoflagellates (Supplementary figure 2), it is plausible that tubulin related regulatory mechanisms are functionally conserved in dinoflagellates. While direct experimental evidence is currently lacking, our study lays the groundwork for testing this hypothesis using a highly promising model system. It also provides a platform to explore how ocean acidification influences phytoplankton motility through non-genomic regulatory pathways.

## Materials and Methods

### Culture maintenance/Cell Culture

Non axenic, uni-algal cultures of *Lingulodinium polyedra* (Previously *Gonyualax polyedra*) (CCMP—1936, kind gift from Prof. David Morse, originally obtained from Provasoli-Guillard National Center for Marine Algae and Microbiota, East Boothbay, ME, USA) was grown in L1 media. The pH of the medium was adjusted to about 8.2±0.2. All cultures were maintained under 12:12 Light: Dark (LD) regimes of cool fluorescent light with intensity of about 60 µmol photons m^−2^·s^−1^ at 18 *±* 1 degree Celsius. *Amphidinium carterae* and *Effrenim voratum* was grown in F/2 sea water media of pH 8±0.2. Cultures were maintained like *L*.*polyedra*.

### Cell density/Cell count measurement

For *L. polyedra* and *E. voratum*, cultures were diluted at a 1:4 ratio (200 µL culture to 800 µL of L1 or F/2 medium). 10 µL of this was mixed with 790 µL of Lugol’s iodine solution for counting. For *A. carterae*, cell counts were performed by directly mixing 20 µL of culture with 980 µL of Lugol’s iodine solution. In all cases, 10 µL of the diluted sample was placed on a glass slide and observed under a Leica DM500 microscope equipped with an ICC50 W camera for cell counting.

### pH Alteration using CO2 and visualization of cellular motility

80 ml of *L. polyedra, A. carterae*, and *E. voratum* cultures with a cell density of about 8000 cells ml^-1^, 0.4 million cells ml^-1^, and 20000 cells ml^-1^, respectively were transferred to 500 ml borosil Glass beakers. The pH of the cultures was measured using either a handheld digital pH meter or a vernier GO direct pH probe calibrated with pH 4.00, 7.00 and 9.00. Occasionally, we cross-calibrated the pH with the Hanna pH probe (HI7698494/4). The beakers were covered with aluminum foil. Using the CO_2_ airflow controller through a gas control box equipped with a regulator, a CO_2_ line using a ⅛ inch teflon tube fitted with a 1 mL tube was inserted into the test group beakers through a minimal hole punched in the aluminum foil. The CO_2_ was not bubbled into the medium, instead it was introduced into the airspace above the medium. The beakers were kept in a shaker with slow shaking and the pH was monitored. Under this condition CO_2_ flow for 30 seconds, 15 seconds, 8 seconds and 2 seconds resulted in pH of 5.8*±*0.1, 6.1*±*0.1, 6.5*±*0.1 and 7.1*±*0.1 respectively. After 10 minutes, 20 to 30***µ***l of the sample and control was observed under Leica microscope and video was captured for 30 seconds. In the case of *E voratum* and *A. carterae* CO2 was delivered with the similar parameters as mentioned above for *L. polyedra*, resulting in a pH of 6.5±0.1 for *E. voratum* and 6.2±0.1 for *A carterae*. After 5 to 10 minutes, 20 μL of the sample was observed under the microscope. At least 3 biological replicates and several technical replicates from each biological replicate were monitored for each of the 3 different species. Triplicates were conducted on three separate days, with each replicate performed at the same time of day to control potential circadian influences on cellular motility. All imaging was performed within the central (bulk) region of the droplet to minimize boundary-related artifacts.

To alter the pH biochemically the L1 medium pH was altered using 1N HCl. The pH was adjusted to around 7±0.1 before adding the cells. Once the cells were added the pH stabilized to 7.1, which was monitored by the handheld digital pH meter. 10 minutes after stabilization of the pH 10 ***µ***l of sample and control cells were visualized and documented as described above.

To revert the pH, fresh L1 media of pH 8.5 was added to the culture with pH 6.5 after 10 minutes. The resultant pH was about 8. Videos were captured before CO_2_ Shock, 10 minutes after pH shock and 16 hours after transfer into fresh media and reverting their pH to 8.

To observe flagellar behavior in cells incubated in an acidified medium, we used an ultrafast frame imaging Thorlabs camera (CS135MUN) mounted on a Leica DMP4 polarizing microscope equipped with a 20X objective. Imaging was conducted at a frame rate of 165 fps. For visualization purposes, representative videos were compiled at 20 fps. All imaging was performed within the central (bulk) region of the droplet to minimize boundary-related artifacts. All experiments were performed in triplicate on three separate days, with each replicate conducted at the same time of day to control for potential circadian variation.

### Nitrogen

80 ml of *L. Polyedra* cultures with respective cell density of about 8000 cells ml^-1^ were transferred to 500 ml beakers and N_2_ gas was applied for 30 Seconds with the flow rate that is 5 times higher than the CO_2_ as described above. Video was captured for 30 seconds before treatment and 10 minutes post-treatment using Leica microscope with 10***µ***l to 30***µ***l of the sample. At least 3 biological replicates and multiple technical replicates from each biological replicate were monitored.

### Video and Data Analysis

For *L. polyedra*, the video was captured in AVI format and then sliced into individual frames as PNG files using Python packages moviepy and CV2. The frames from each video were further used to analyze from multiparticle tracking package Trackpy-0.6.2 [11] based on the Crocker and Grier algorithm [12]. The following parameters were used in Trackpy to identify and link the cells: memory=99, size =67, separation=15 pixels and search range=25. The ‘search range’ defines the maximum distance a cell can move between frames, while ‘memory’ specifies the maximum number of frames a cell can disappear before being assigned a new identity. For A. carterae and E. voratum, all experimental conditions were kept identical except for the parameter values, which were modified as specified. *E. Voratum*: Size(diameter) =55, memory =66, separation =15, search range= 25 and *A. carterae*: Size(diameter)= 47, memory= 66, separation= 15, search range= 25. The speed data was computed from the time series of (x, y) coordinates corresponding to the cell’s centroid using the formula of speed=euclidean distance/time. Since cells might disappear and reappear between frames, leading to discrepancies in frame numbers, these variations were accounted for in the Python script during the speed calculation. The resulting speed data was then visualized using violin plots created with the Matplotlib and Seaborn libraries in Python.

## Acknowledgement

This study was supported by the Ministry of Earth Science, Government of India (MoES grant MoES/36/OOIS/Extra/80/2020) and Department of Biotechnology, Government of India (DBT grant BT/PR32511/BRB/10/1803/2019), SR acknowledges the Annual Research Grant from Ashoka University. AK was supported from the MoES manpower grant and SRT was supported by a PhD fellowship from Ashoka University. We acknowledge Dr. Pramoda Kumar and Ankit Gupta for generously allowing us to use their polarized microscope imaging system and for their valuable assistance during the imaging experiments.

## Author contributions

**AK:** Conceptualization of experiments; data curation; software; experimentation - designing and performing; formal analysis; validation; investigation; visualization; methodology; writing – review and editing. **SRT:** Standardization of cell tracking using trackpy, help AK to generate the cell trajectories and some formal analysis, polarized microscopy imaging and their analysis, writing – review and editing **AS:** performed some experiments the cell motility under normal and pH shock. **SR:** Conceptualization: conceiving the original idea; experiment designing; resources; supervision; project administration; funding acquisition; writing – original draft; writing – review and editing.

## Data Availability

The code will be available upon request directed to the corresponding author.

