## Supplementary Figures for "pH induced motility pattern change in a marine dinoflagellate"

**S1A**

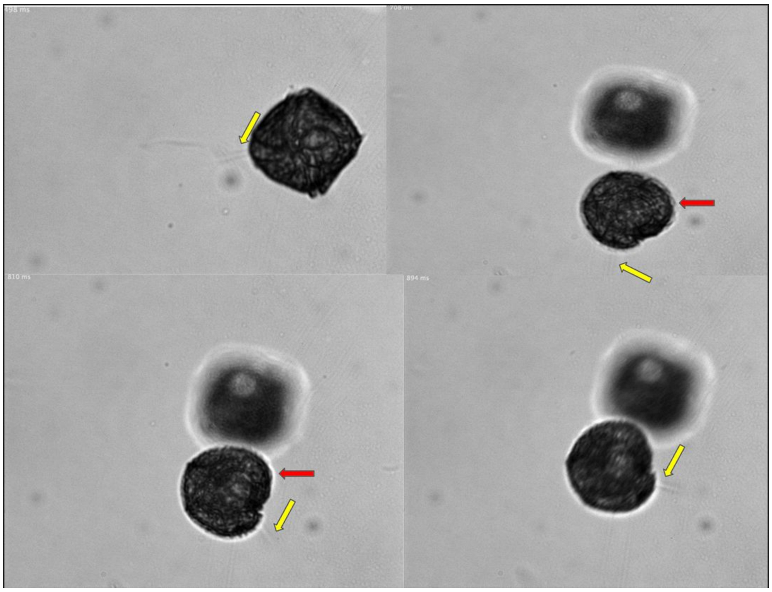

**S1B**

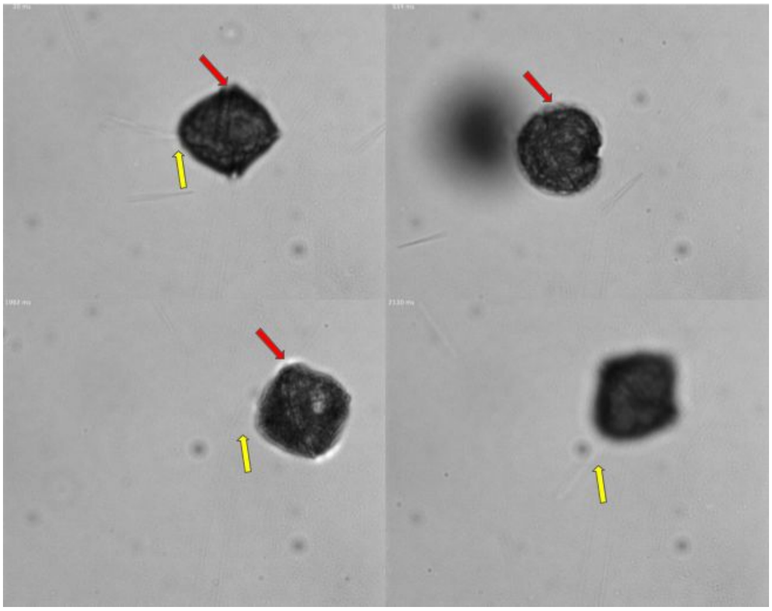

**Supplementary Figure 1: Flagellar imaging from spirally moving single *L. polyedra* cells under acidicconditions.** The images are processed from the video captured files using the Thorlabs camera with fast frame capture of 165 fps. The images presented here have not been modified in any way, except for cropping and resizing. S1A and S1B are two independent *L. polyedra* cells capture under two experiments on separate days from two separately grown cultures. These are considered as biological replicates.

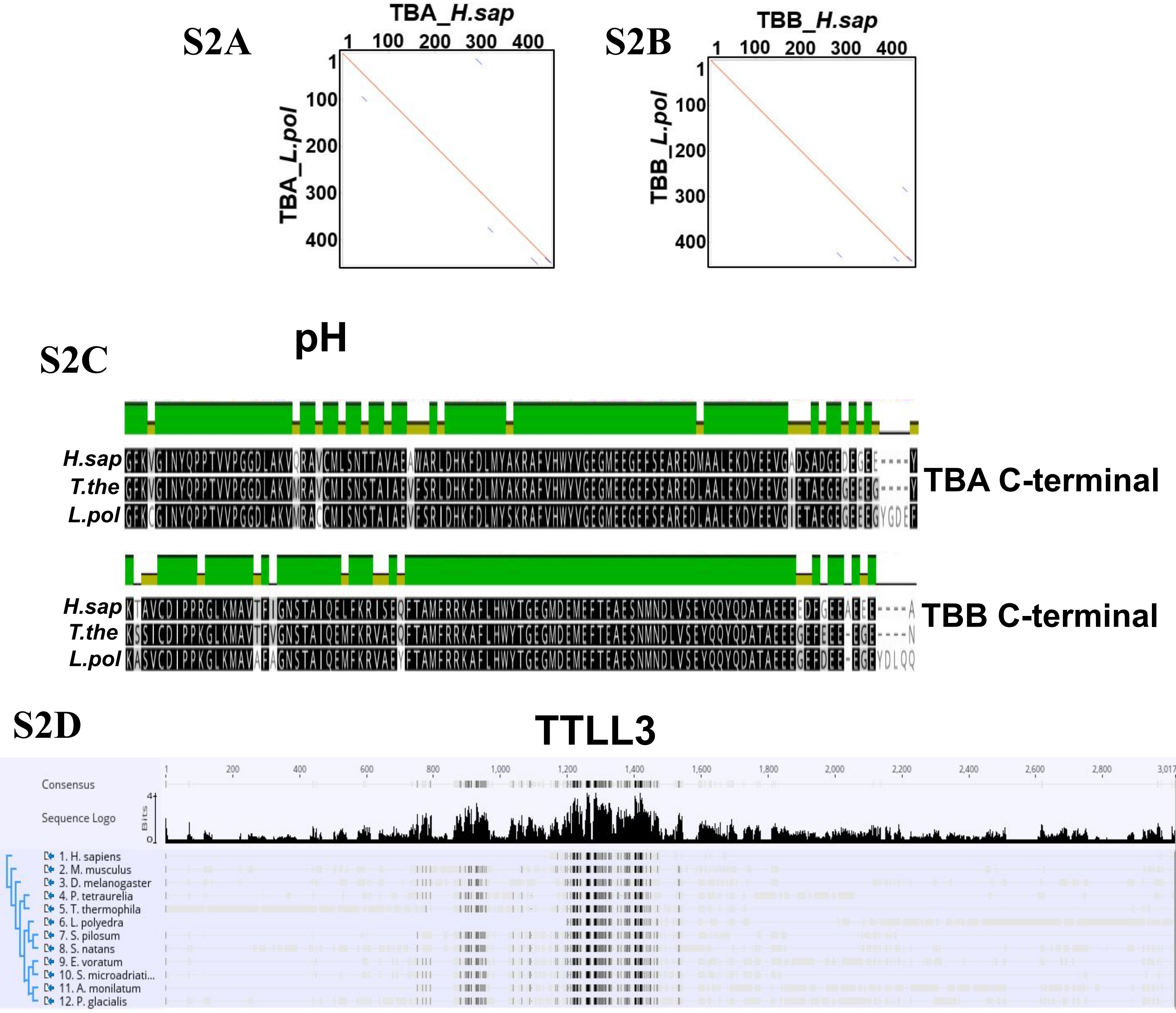


**Supplementary Figure 2: The conservation of tubulin and its modifying enzyme from dinoflagellates to humans:** S2A: Left Panel panel: Dot plot of the sequence alignment of α-tubulin from *L. polyedra* (HBOU01099056.1) and human (NP_116093). Right panel: Dot plot of the sequence alignment of β-tubulin from *L. polyedra* (HBOU01108412.1) and human (NP_821133). The respective *L. polyedra* α and β tubulin sequences were downloaded from the NCBI. S2B: The sequence alignment between the *L. polyedra, T. thermophilia* (Accession P41351.1 and P41352.1) and human C-terminal end (from 350 to the C- terminal end) of the α- and β tubulin respectively. The alignment was generated in geneious software. **S**2C: The phylogenetic tree using sequences of tubulin tyrosine Ligase Like 3 (TTLL3) from mammals, insects, ciliates and dinoflagellates. Using the Geneious prime embedded Muscle multiple sequence alignment followed by RaXML was used for generating the tree. The *T. thermophilia* TTLL3 sequence was used as a query to search for TTLL3 sequences in mammals, insects, ciliates and dinoflagellates. The TTLL3 sequences were downloaded from NCBI and had the following accession AAH98361.1, NP_598684.4, NP_609069.1, XP_001450145.1, XP_001013865.1, HBOU01021278.1, CAE7690824.1, CAE6937418.1, CAJ1436579.1, CAE7392100.1, HBNR01028271, CAK0860068.1.
